# A multiscale cytoskeletal network model for shear rheological property and its evolutionary mechanism

**DOI:** 10.64898/2026.07.13.738349

**Authors:** Han-Lin Liu, Neng-Hui Zhang, Jing-Jie You, Qi-Qi Li, Cheng-Yin Zhang

## Abstract

The cytoskeleton is a dynamic biopolymer network whose shear rheological properties are crucial for cellular physiology and pathology. However, its mechanical behavior spans multiple spatiotemporal scales, and the coupling of dynamic remodeling and viscoelastic dissipation mechanisms poses a challenge for traditional models to comprehensively capture complex cellular responses. This study aims to establish a multiscale cytoskeletal network model that integrates the bio-chemo-mechanical properties of local linked proteins, the viscoelasticity of actin filaments, and their deformation states. Developing a boundary-modified finite element method with an incremental iterative algorithm, we demonstrated the dynamic remodeling of network and the resultant rheological properties of cytoskeleton by extending the predictive time scale to one thousand seconds. The results not only reproduced the short- and intermediate-term power-law creep behavior and long-term strain plateau response of the cytoskeletal network observed in shear rheological experiments, but also indicate that the synergy among the chemo-mechanical coupling of cross-linked proteins and the bending-to-tension transition of actin filaments govern both the network remodeling and its power-law response evolutionary, whereas the steady-state properties of actin filaments determine the long-term network behavior. Simulations of cancerous and drug effects show that cancer-induced softening and reduced filament viscosity lead to accelerated cytoskeletal responses and decreased apparent shear modulus, respectively; and drug-enhanced filament prestress, along with promoting association or inhibiting dissociation of cross-linked proteins, can effectively increase the steady-state shear modulus. These findings advance the understanding of the spatiotemporal evolution and pathological mechanisms of cellular mechanical responses and provide insights for regulating polymer network performance.

## 1 Introduction

The cytoskeleton is a highly structured, dynamic biopolymer network that maintains cell morphology and drives cell movement (Wei et al. 2021). It primarily consists of actin filaments, intermediate filaments, and microtubules (Huber et al. 2013). Through rapid remodeling in response to external or internal changes, it enables cells to migrate, divide, and maintain shape (Liang et al. 2023). Experimental studies have shown that aging and disease significantly alter the cellular elastic modulus, a mechanical response related to cytoskeletal structure and its components such as stress fibers and binding proteins (Kumar et al. 2006; Calzado-Martín et al. 2016; Rianna et al. 2017). However, the intriguing relationships between these microscale components and their emergent macroscale mechanical responses remain unclear. A deeper understanding of cytoskeletal network deformation characteristics and their regulatory mechanisms is therefore crucial for elucidating cytoskeleton-related physiological functions and pathological mechanisms (Liang et al. 2023). This paper focuses on the shear rheological properties of cytoskeleton and will introduce the related research methods and findings in the following sections.

Experimental exploring across multiple temporal and spatial scales has been foundational to the current understanding. Early studies collected the structural characteristics of cytoskeleton using electron microscopy (EM) and fluorescence microscopy (FM) (Burla et al. 2019). Guided by these structural observations, the reconstituted biopolymer networks were developed in vitro to mimic cytoskeletal functions (Foffano et al. 2016; Burla et al. 2019). Studies on the shear behavior of these reconstituted networks can be generally summarized from two aspects: statics (Storm et al. 2005; Fletcher and Mullins 2010) and dynamics (Gardel et al. 2006; Lieleg et al. 2010; Müller et al. 2014). In terms of statics, shear experiments have shown that the cytoskeleton exhibits significant nonlinear elastic behaviors, particularly as the external load increases, where its overall stiffness undergoes orders-of-magnitude increment (Storm et al. 2005). Combined prior single-molecule mechanical experiments and recent structural observations (Storm et al. 2005; Hatami-Marbini and Rohanifar 2021), some scholars ascribed this nonlinear elastic behavior to the process of network structure transition from bending- to tension-dominated fiber deformation. In terms of dynamics, rheological experiments further found that the cytoskeleton exhibits multi-power-law behaviors across different time/frequency regimes, similar to that of whole cells (Gardel et al. 2006; Lieleg et al. 2010; Kumar et al. 2013). By coupling mechanical tests with high-resolution structural measurements, some scholars associated this rheological behavior with force-dependent dissociation/association of cross-linkers (Lieleg et al. 2010; Huang et al. 2017), configurational transitions of local filaments (Risca et al. 2012), and remodeling of cytoskeletal network (Majumdar et al. 2018). Other microscopic rheological mechanisms, such as the number of filaments (Gavara et al. 2016) and the disentanglement and slippage of physical entanglements between filaments, have been also identified (Kurniawan et al. 2012).

Modeling the cytoskeleton by integrating mechanical, chemical, and biological theories can reproduce experimental phenomena and reveal the underlying mechanisms behind them. Scholars have established different cytoskeletal models at the relevant spatiotemporal scales (Yamaoka et al. 2012; Burla et al. 2019). These models can be generally categorized into continuum models and discrete structural models (Rodriguez et al. 2013; Wang et al. 2021). Continuum models treat the cytoskeleton as continuum materials and employ constitutive models with different levels of complexity to describe its mechanical behavior based on different experimental phenomena. Single-phase continuum models can reproduce the mechanical response of the cytoskeleton observed in macroscopic shear experiments to some extent (Trepat et al. 2007); however, the homogenizing treatment of the whole cytoskeleton is difficult to accurately describe the local non-uniformity, and sometimes may ignore the contribution of certain components (Wang et al. 2021). Multi-phase continuum models can describe the interactions among different cytoskeletal components and partially reflect the viscoelastic physical mechanisms (Fallqvist and Kroon 2013; Moeendarbary et al. 2013); yet they are still hard to capture the dynamic evolution of cytoskeletal response to the localized non-uniform interactions (Rodriguez et al. 2013; Wang et al. 2021). Discrete structural models view the cytoskeleton as a network composed of discrete elements, and they can be further divided into specific framework models based on structural engineering and discrete network models based on statistical materials science (Burla et al. 2019). Specific framework models, by employing preset self-equilibrating structures, can qualitatively reveal the distribution of prestress, force transmission, and their effects on the overall properties of the cytoskeleton (Xu et al. 2018; Palumbo et al. 2022); however, the assumed structural forms and material parameters are difficult to reflect the dynamic evolution of cross-linkers and network remodeling. Discrete network models, by defining material properties and cross-linked rules, can emerge behaviors such as nonlinear stiffening and network remodeling from the interactions of the components (Burla et al. 2019); however, these models focus on revealing specific mechanisms (Kim et al. 2015) by often simplifying fibers and cross-linkers as elastic elements (Zeng et al. 2012), and this simplification make it difficult to fully incorporate the time-dependent properties of components and the dynamic evolution of cross-linkers into the overall rheological behavior of the cytoskeleton (Burla et al. 2019).

In pursuit of realism and effectiveness, discrete structural network models have been continuously developed. Specifically, a major research focus has been on how conformational changes in actin filaments, driven by thermal fluctuations, influence the overall network response. Scholars have introduced the effects of thermal fluctuations on filament configuration by assigning pre-curved filament shapes (Onck et al. 2005) or using dynamical approaches (Lin et al. 2014). By constructing finite element (FE) network models based on discrete filaments, they have studied the mechanical response of cytoskeletal network and confirmed that the stiffening of the cytoskeleton is closely related to the network structure and the transition of filament deformation (from bending- to tension-dominated states). Another widely investigated aspect is how the dynamic evolution of cross-linkers affects the network-wide response. By simplifying cross-linkers into elastic spring (Zagar et al. 2015), and introducing the dissociation/association (Broedersz et al. 2014) into FE network models using the modified Bell model (Bell 1978), scholars have demonstrated that the cross-linked network behaves as a solid at timescales shorter than the characteristic dissociation/association time (less than half a second) but exhibits fluid-like behavior at longer timescales (several minutes). Additionally, by establishing Brown/FE-Langevin dynamics model incorporating cross-linking kinetics (Wei et al. 2021; Maxian et al. 2022), scholars have demonstrated the influence of filament prestress, deformation transition, and cross-linker properties on the network remodeling and the bulk performance of cytoskeleton; however, constrained by computational limitations (Saunders and Voth 2013), the predictive capability for mechanical responses is currently limited to shorter timescales (≤ 60 s).

This study aims to provide a cytoskeletal network model by establishing cross-scale correlations among bio-chemo-mechanical properties of local cross-linkers, viscoelasticity and deformation transition of single actin filament, and dynamic remodeling and rheological properties of bulk cytoskeleton. First, inspired by micromechanical concepts, we selected a cytoskeleton network composed of finite quadrilateral cellular elements as the representative unit (Lin et al. 2014; Abhilash et al. 2014), based on structural features observed in EM images (Palmer and Boyce 2008). Specifically, drawing from experimental observations of single-scale cytoskeletal components (Kumar et al. 2006; Huang et al. 2017), we described the viscoelastic properties of actin filaments using a three-parameter solid model (Lyu et al. 2020). We also simplified the chemo-mechanical coupling effect of cross-linkers as rotational springs using the Bell model (Bell 1978; Nam et al. 2016); and simplified physically entangled nodes as hinges, maintaining a statistical proportional relationship with cross-linkers (Kurniawan et al. 2012). As a result, a multiscale non-uniform network model of the cytoskeleton was established based on experimental statistical patterns (Kurniawan et al. 2012). Second, we analyzed the forces and deformations of actin filaments at the boundaries and modified the boundary terms in the network model by using the microelement method; and developed an incremental iterative FE method with a time-discrete nested format to incorporate and dynamically update the time-dependent mechanical properties of actin filaments and the bio-chemo-mechanical characteristics of cross-linkers. Furthermore, by solving the displacements of network nodes, we statistically aggregated the deformation of all cytoskeletal elements to characterize the bulk deformation features and shear rheological properties of the network across different time scales. Finally, considering the cancerous and drug effects on cytoskeletal structure and multiple components, we explored their influence on the apparent shear modulus by controlling relevant component and structural parameters. Our simulations not only reproduced the shear rheological response of the cytoskeleton across different time scales, but also explored the effects of cancerous/drug-related parameters on the bulk mechanical properties. These insights provide a mechanistic explanation and theoretical foundation for inducing cellular network structural changes and for developing polymer network materials.

## 2 Materials and methods

Based on existing experimental observations (Fig. 1(a)), we will establish a multiscale FE model to efficiently and rationally predict the shear rheological response of cytoskeletal network. First, the cytoskeleton will be simplified as a network of viscoelastic actin filaments and dynamically linked nodes (Figs. 1(b) and 1(c)). Second, a numerical model incorporating both network remodeling and filament viscoelasticity will be established to simulate shear rheological experiments. Finally, structural and material parameters will be determined using existing experimental and simulated data for single-scale studies, ensuring experimental grounding and model verifiability.

**Fig. 1.**
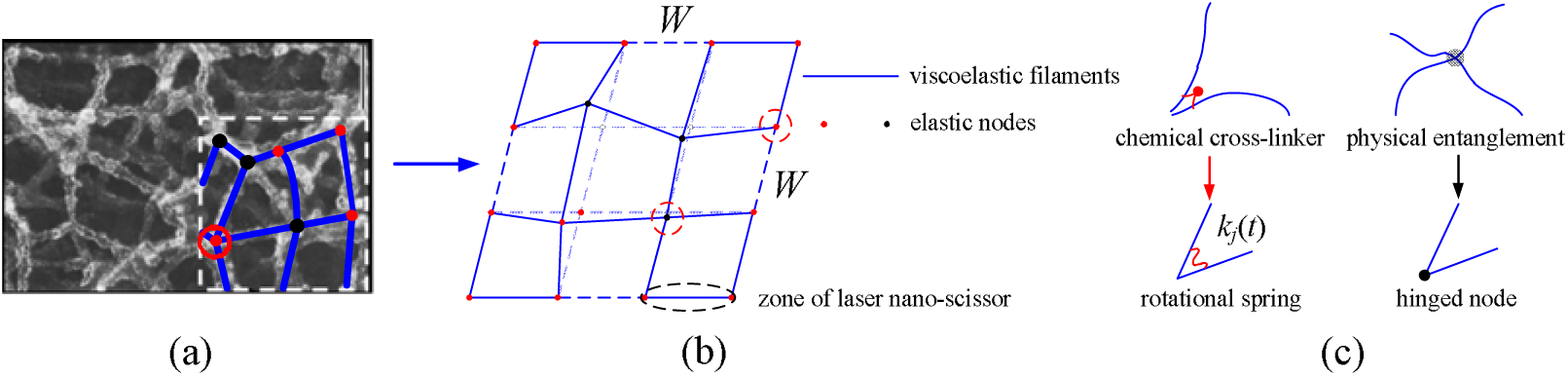
Schematic diagram of multiscale cytoskeletal network model: (a) EM image, reprinted from Acta Biomaterialia, Vol 4(3), Jeffrey S. Palmer, Mary C. Boyce, Constitutive modeling of the stress-strain behavior of F-actin filament networks, Pages 597–612, Copyright (2008), with permission from Elsevier, (b) network model, (c) node model.

### 2.1 Composition of cytoskeletal network and its simplification

To ensure that our simplified cytoskeletal network models can capture key physical mechanisms while remaining computationally feasible, we developed multiscale models based on EM, FM, and atomic force microscopy observations (Huber et al. 2013; Burla et al. 2019; McArthur et al. 2025).

The cytoskeleton is widely recognized as a dynamic biopolymer network (Wei et al. 2021). Therefore, at the network scale, we employ a composite mechanics approach, representing the cytoskeleton with a two-dimensional network composed of finite quadrilateral cellular elements (Fig. 1(b)) (Gardel et al. 2006; Palmer and Boyce 2008). This unit characterizes uniformly distributed cytoskeletal filaments within a three-dimensional space (*W*×*W*×2*r*), with its volume fraction *f*_V_ given as (Zhang et al. 2021)

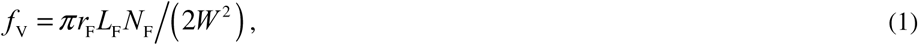

where *r*_F_, *L*_F_, and *N*_F_ represent the radius, length, and number of actin filaments, respectively, and *W* denotes the edge length of the simulation domain.

Experiments demonstrate that cytoskeletal stress fiber (a specific type of filament) exhibits time-dependent mechanical properties (Kumar et al. 2006). Therefore, at the filament scale, we model the filaments as prestressed three-parameter viscoelastic solid rods with the constitutive relationship given as (Christensen, 1982; Zhang and Xing 2008)

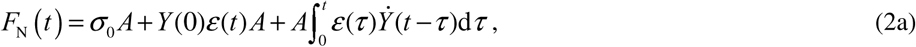

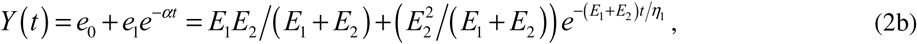

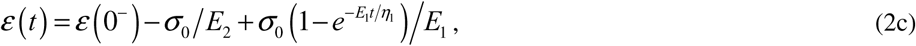

where *F*_N_, σ_0_ and *A* are the axial force, prestress and cross-sectional area of filament, respectively; *Y*(*t*) is the time-dependent relaxation modulus, ε(*t*) is the deformation accounting for the history effect ε(0^−^). *E*_1_ and η1 are the spring stiffness and viscous coefficient of the Kelvin solid element, respectively, while *E*_2_ is the stiffness of the series spring element.

At the node scale, based on experimentally derived bio-chemo-mechanical properties, the researchers often classified the linked nodes (Fig. 1(c)) into two types: physically entangled nodes connected by weak interactions, and chemically cross-linked nodes connected by specific cross-linkers (Kurniawan et al. 2012; Broedersz et al. 2014). Physically entangled nodes can be simplified as hinge nodes, and are assumed not to fail (Kurniawan et al. 2012); chemically cross-linked nodes can be modeled as rotational springs with force-regulated dissociation/association rates (Broedersz et al. 2014; Kurniawan et al. 2012), following the Bell model (Bell, 1978). The evolution of the cross-linkers binding density *N*_b_ is given by (Nam et al. 2016)

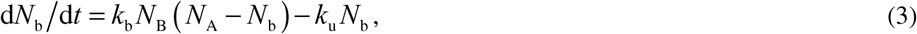

where *N*_A_ and *N*_B_ are the ligand density and receptor density, respectively (Nam et al. 2016); *k*_u_ and *k*_b_*N*_B_ are the force-regulated dissociation and re-association rates, respectively (Suppl. Mat. B.). In addition, the count of entangled nodes *R*_e_ and cross-linked nodes *R*_cl_ follow the statistical relationship, *R*_e_ = *s R*_cl_, where *s* is the proportionality constant, based on FM experimental observations (Kurniawan et al. 2012).

### 2.2 Numerical model of cytoskeletal network

To simulate the shear rheological response, we will employ the FE method to couple the multiple components. The process involves: generating a random statistical network based on experimental patterns and assembling the overall stiffness matrix, applying boundary constraints and equivalent nodal loads, and solving the numerical model using a time-discretization scheme combined with nested Newton-Raphson iteration.

#### 2.2.1 Generation of overall stiffness matrix

We constructed a random statistical network representing a non-uniform cytoskeleton by first introducing initial angles and random displacements to the internal nodes of a uniform network (Figs. 2(a) and 2(b)), and then assigning hinge or rotational spring properties to these nodes based on experimental statistics (Kurniawan et al. 2012) (Fig. 2(c)).

**Fig. 2.**
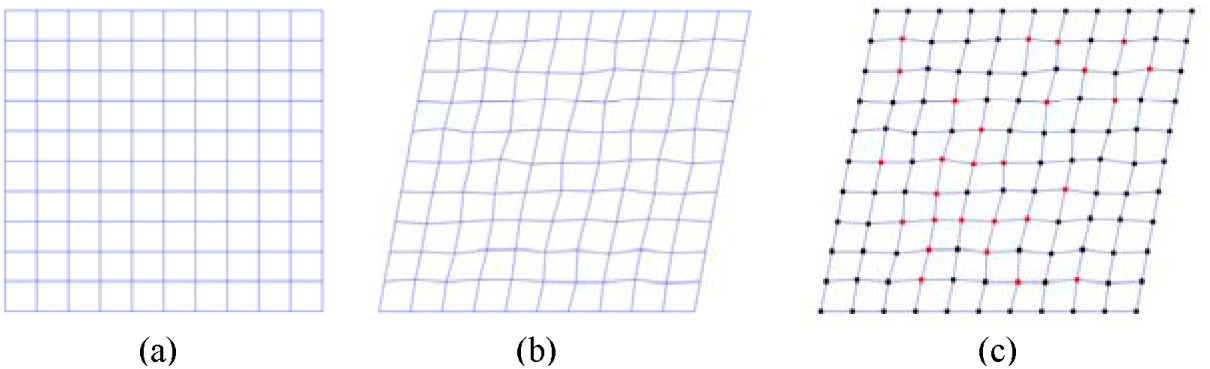
Generation of non-uniform random cytoskeletal networks: (a) uniform network, (b) introduction of initial angles and random node displacements, (c) proportional assignment of internal node types.

Structural discretization and encoding the non-uniform random cytoskeletal network enable the localization of its nodes and units. Considering the viscoelastic properties of filaments, the local constraints imposed by nodes, and the angle between filaments and the global coordinate system, the element stiffness matrix of filaments in the global *XOY* coordinate system is (Suppl. Mat. A.):

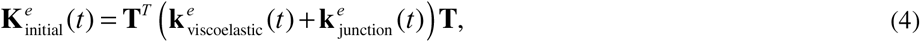

where **k***^e^*_viscoelastic_(*t*) is the local stiffness matrix of viscoelastic filaments, influenced by the time-dependent relaxation modulus *Y*(*t*) and geometric parameters; **k***^e^*_junction_(*t*) is the stiffness contribution from the linked nodes to the local element matrix, affected by the dynamic bio-chemo-mechanical properties of nodes; **T** is the coordinate transformation matrix. And the overall stiffness matrix is assembled by superimposing all element matrices **K***^e^* (*t*).

#### 2.2.2 Boundary handling and load generation

Numerical simulations have shown that under an applied shear stress, representative units within the cytoskeleton experience boundary constraints and loading analogous to those on the cell surface (Gardel et al. 2006; Moeendarbary et al. 2013). Regarding the load-induced restriction by intracellular fluid, the lateral boundary conditions of the cytoskeletal network can be approximately handled by considering the unit shearing stress τ*H*(*t*), where τ = τ_0_/*W* (Gómez-Gonzále et al. 2020). To simulate this, we applied fixed constraints to the lower boundary of the cytoskeletal network to represent the basal constraints. Meanwhile, a uniformly distributed force τ*H*(*t*)*A*_c_, representing the resultant force of the bulk cytoskeletal shear stress, is applied to the upper and lateral boundaries to model the step shear loading, where *A*_c_ is the axial cross-section area of actin filaments. These loads are applied to the boundary elements *e* of cytoskeletal network and can be equivalently treated as fixed-end forces (Suppl. Mat. A.).

Since cytoskeletal filaments at the outer boundary are subjected to eccentric surface forces, additional boundary constraints should be considered to enhance the predictive accuracy of FE model. Using the infinitesimal element method, we analyzed the boundary elements *e* and determined the following quasi-static governing equation (Suppl. Mat. C.)

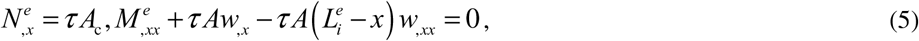

where *N^e^*and *M^e^* are the axial force and bending moment of filament element *e* at the current time step, respectively.

At the current time step, the internal nodes (subscript I) and boundary nodes (subscript B), together with the overall stiffness matrix, displacement vector, and load vector, are represented in block form as

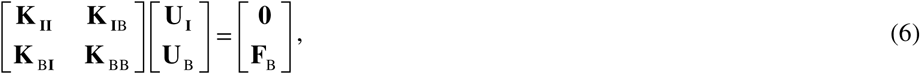

where **K**_II_ is the sub-stiffness matrix for elements with both ends as internal nodes; **K**_IB_ and **K**_BI_ are the coupling sub-stiffness matrices between internal and boundary node degrees of freedom; and **K**_BB_ is the sub-stiffness matrix for elements with both ends as boundary nodes, which requires geometric correction (Suppl. Mat. C.). **U**_I_ and **U**_B_ are the displacements of internal and boundary nodes, respectively, and **F**_B_ is the force applied to the boundary nodes.

#### 2.2.3 Numerical solution methods

Considering the transition in local cytoskeletal filament deformation (between bending-and tension-dominated states) caused by node bio-chemo-mechanical characteristics, filament viscoelasticity and force transmission, we employed a nested solution structure for quasi-static analysis and displacement determination of the cytoskeletal network. This is achieved through time discretization combined with an incremental iterative method.

In the time domain, to efficiently capture the responses from short-term (10^–3^ s) to long-term (10^3^ s) scales, a segmented adaptive time-stepping strategy on a logarithmic scale is adopted; and in the space domain, the Newton-Raphson iterative method is employed until convergence is achieved at each time step. The collaborative solution steps are as follows (Suppl. Mat. A.).

**Step 1.** Self-balance of initial state. For *t* = 0^−^, the prestressed cytoskeleton exhibits self-balancing properties (Kumar et al. 2006; Wei et al. 2021). And its initial physical equilibrium state can be numerically determined when the residual norm for all nodes satisfies //**R**_0_^(*k*)^// ≤ 10^−6^ (Iravani et al. 2020).

**Step 2.** Quasi-static equilibrium process. For *t* ≥ 0^+^, the initial displacement condition is taken from the previous time step. Considering the viscoelastic deformation of filaments and node constraints, internal nodes are assumed to be in approximate equilibrium at each time step. Thus, at current iteration *k*, both the residual norm conditions (//**R**^(*k*)^// ≤ 10^−6^) and the displacement convergence conditions (//**U**^(*k*)^// ≤ 10^−4^) should be satisfied (Iravani et al. 2020). Once the displacement correction **U**^(*i*+1)^ = **U**^(*i*)^ +Δ**U**^(*i*)^ meets these conditions, the converged displacement **U**_(*n*+1)_ is saved.

**Step 3.** Judgment of cross-linked node states. Based on Eq. (3), the node force, the dissociation and re-association rates (*k*_u_ and *k*_b_*N*_B_) are updated under the current deformation. Node state transitions are then evaluated by

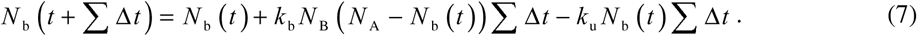

If |*N*_b_(*t* + ΣΔ*t*)–*N*_b_(*t*)| ≥1/*R*_cl_, where *R*_cl_ is the count of cross-linked nodes, an integer jump (i.e., a cross-linker dissociation) is detected. It means that, one of the cross-linked nodes transitions to a dissociated state. The rotational stiffness of that node is changed, and the element stiffness matrix is reset and then embedded into the overall stiffness matrix to reflect this change. **Step 2** is repeated until the count of cross-linked nodes reaches a dynamic equilibrium state, i.e. |*k*_b_*N*_B_(*N*_A_–*N*_b_) –*k*_u_*N*_b_| <1%, after which the results at the current time step are output.

**Step 4.** Post-processing and result output. After obtaining the quasi-static solution for the cytoskeletal network at each time step, the shear strain creep response and apparent shear modulus of the cytoskeletal network are given by (Efremov et al. 2020; Iravani et al. 2020)

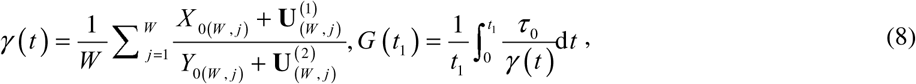

where *j* is the column index of the cytoskeletal network, *X*_0(*W*_, *j*) and *Y*_0(*W*_, *j*) are the initial (*t_n_* = 0) global coordinates of the *j*-th boundary node; τ0 is the amplitude of unit shear stress, and *G* (*t*_1_) is the apparent shear modulus at time *t*_1_. To reveal microstructural evolution, the deformation states of all filaments are statistically classified as bending- and tension-dominated modes, with proportions *p*_b_(*t*_1_) and *p*_s_(*t*_1_), respectively. To describe the bulk configuration change, the Shannon entropy is calculated as (Sajjadi et al. 2024)

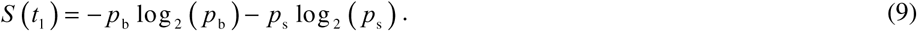

For the calculation of network deformation, the buckling state of all filaments is assessed at each time step using the critical buckling load formula (Yu et al. 2025). After conducting mesh independence verification, we selected a cytoskeletal network composed of 11×11 units for simulation. During the calculation process of global network, we observed the changes in results by reducing Δ*t* to verify the stability of the computational results and to determine whether to output them, ensuring the independence of the time step (Suppl. Figure S1).

### 2.3 Simulation of shear rheological experiments

#### 2.3.1 Selection of model parameters

To reproduce the creep response characteristic of the cytoskeleton experiments (Huber et al. 2013; Burla et al. 2019; Chang et al. 2024) and to further investigate the effects of cancer-related or drug-modulated parameters on its apparent shear modulus (Hayashi and Iwata 2015; Rigato et al. 2017; Linklater et al. 2021), appropriate model parameters should be selected. These include structural parameters (filament volume fraction *f*_V_, Eq. (1)), filament properties (viscoelastic tri-parameters *E*_1_, η1, *E*_2_, and prestress σ0, Eq. (2)), and the bio-chemo-mechanical properties of cross-linkers (Eq. (4)). Due to the lack of direct parameter data, the material and structural parameters for the multiple components are integrated from existing experimental measurements and numerical simulations. The structural parameters of the network geometry and fiber/filament dimensions are listed in Table 1, based on data from laser nanoscissor experiments, molecular dynamics, and FE simulations (Kumar et al. 2006; Huber et al. 2013).

**Table 1.** Structural parameters of cytoskeletal networks from experiments/simulations (Kumar et al. 2006; Huber et al. 2013)

| Physical variables | Range of variation | Values used in our study |
| --- | --- | --- |
| Length of stress fiber $L_s$ ($\mu\text{m}$) | 6.64 | 6.64 |
| Radius of stress fiber $r_s$ (nm) | 150 – 800 | 200 |
| Length of filament $L_F$ ($\mu\text{m}$) | 0.8 – 3 | 1 |
| Radius of filament $r_F$ (nm) | 7 – 300 | 70 |

The data on viscoelastic properties of cytoskeletal filament is scarce. Here, we used laser nano-scissor experimental data on stress fibers from Kumar et al. (2006) to determine the prestress and viscoelastic parameters for our model of cytoskeletal filaments. The fitting procedure involved simplifying the experiment as a prestress unloading problem (Suppl. Mat. B.), and the resulting parameters sets for different drug treatments are listed in Table 2.

**Table 2.** Prestress and viscoelastic parameters of actin filaments under drug treatments.

| Drug | $\sigma_0$ (Pa) | $\eta_1$ (Pa·s) | $E_1$ (Pa) | $E_2$ (Pa) | Goodness of fit |
| --- | --- | --- | --- | --- | --- |
| Non | 179.71 | 6278.23 | 655.95 | 395.33 | 0.983 |
| ROCK | 39.98 | 4257.14 | 367.91 | 151.99 | 0.970 |
| MLCK | 1.99 | 241.38 | 149.54 | 188.20 | 0.766 |

**Table 3.** Bio-chemical parameters of chemically cross-linked nodes (Wei et al. 2021)

| Variables and physical meaning | Range of variation | Values used in our study |
| --- | --- | --- |
| Dissociation rate $k_u^0$ (s<sup>-1</sup>) | 0.09 – 15 | 15 |
| Association rate $k_b^0 N_B$ (s<sup>-1</sup>) | 0.1 – 60 | 60 |

For cross-linkers, the intrinsic dissociation and association rates (*k*_u_^0^ and *k* ^0^*N* ) under zero loads were taken from the literature (Nam et al. 2016; Wei et al. 2021), as listed in Table 3. And the rotational spring stiffness *k*_0_, representing the mechanical constraint of a cross-linker, was described by an empirical function of the filament volume fraction *f*_V_ established by Zhang et al. (2021), *k*_0_ = *C*_1_ + *C f ^C^*3, where *C*_1_ = 1.07×10^−5^ N·μm/rad, *C*_2_ = 7.13×10^−2^ N·μm/rad, and *C*_3_ = 3.51.

#### 2.3.2 Simulation for cells shear rheological experiments

After determining the model parameters, we conducted a series of simulations: i) to predict the cytoskeletal creep response under step shear stress, ii) to analyze the underlying microscopic mechanisms, iii) to simulate the effects of cancerous transformation, and iv) to evaluate the effects of drug-treated parameters on the apparent shear modulus.

##### Model validation

First, we validated our model by considering normal cytoskeletal networks with different filament volume fractions (specifically, 26.4%, 30.8%, and 35.2%). Then, we simulated their macroscopic creep response under a step shear stress of 0.03 Pa and compared the results with the experimental data of Gardel et al. (2006), as shown in Fig. 3.

**Fig. 3.**
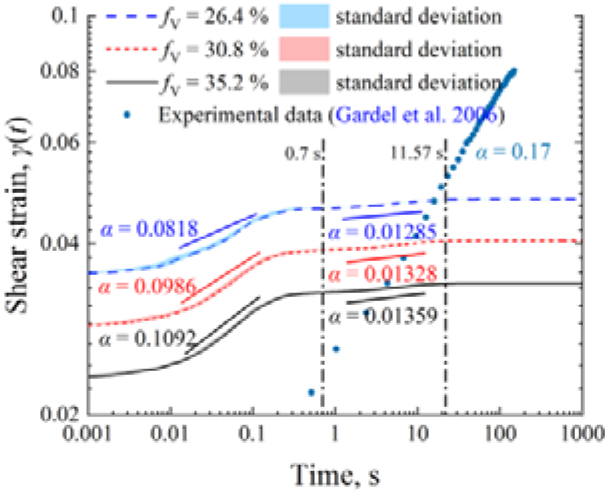
Creep response of the cytoskeletal networks with varying filament volume fractions under step shear stress.

##### Analysis of microscopic mechanisms

To elucidate the underlying microscopic mechanisms, we analyzed the effects of single filament deformation and node bio-chemo-mechanical transitions on bulk network remodeling and power-law rheology. This was done for a drug-free cytoskeleton network with a filament volume fraction of 35.2% (parameters from Table 2) and a proportionality constant of 0.5 between entangled and cross-linked nodes (Kurniawan et al. 2012). The results, showing the evolution of shear strain, network entropy, filament deformation modes, and cross-linker states, are presented in Fig. 4.

**Fig. 4.**
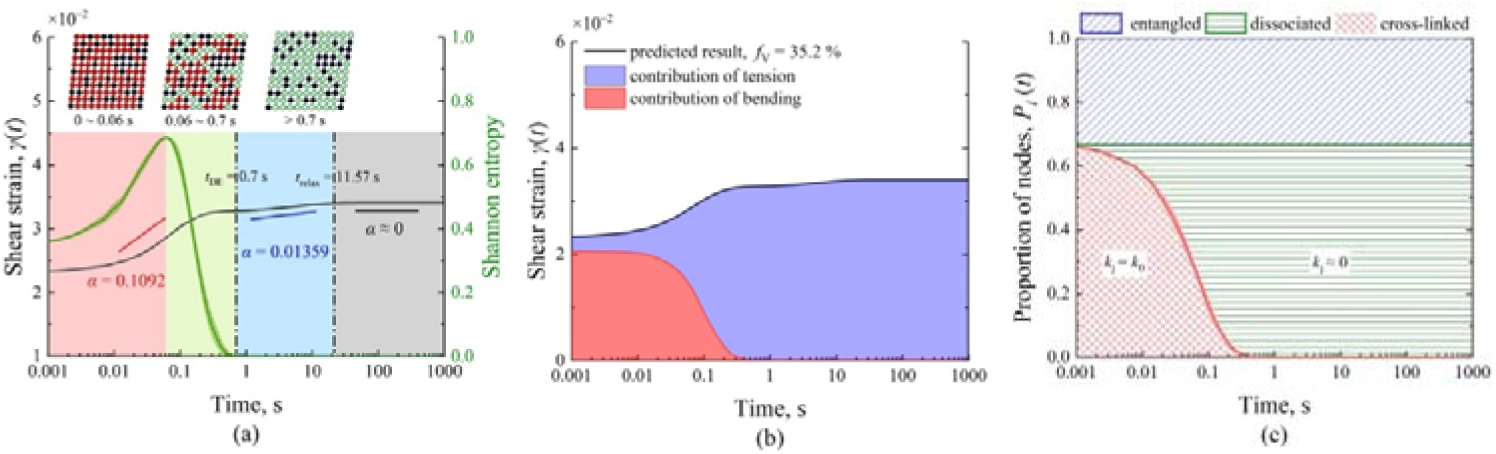
Response evolution of the cytoskeletal network under step shear stress (filament volume fraction 35.2%), (a) bulk shear response and network remodeling (solid line represents shear strain, dashed line represents Shannon entropy), (b) filament deformation contribution, and (c) local node dynamic bio-chemo-mechanical property transition.

##### Simulating cancerous softening

To investigate the impact of cancerous transformation, we simulated moderate- and considerable-softening cancer-like cytoskeletal networks based on the normal network (*f*_V_ = 35.2%), adopting Chen et al. (2023) as an operational definition of parameterized softening levels rather than clinical stages. The normal state parameters were calibrated from the literature (Suppl. Mat. B.), and cancerous changes were selected within reported ranges (Wu et al. 2022; Dessard et al. 2024). And the key parameters were adjusted relative to the normal state: the filament prestress was reduced by 77.8% or 98.9%, viscosity coefficients by 32.1% or 96.2%, density by 12.5% or 25%; and the cross-linker dissociation rate was increased by 100% or 300% (2023). The apparent modulus was calculated using Eq. (8) for normal, moderate-softening, and considerable-softening states, with results shown in Fig. 5.

**Fig. 5.**
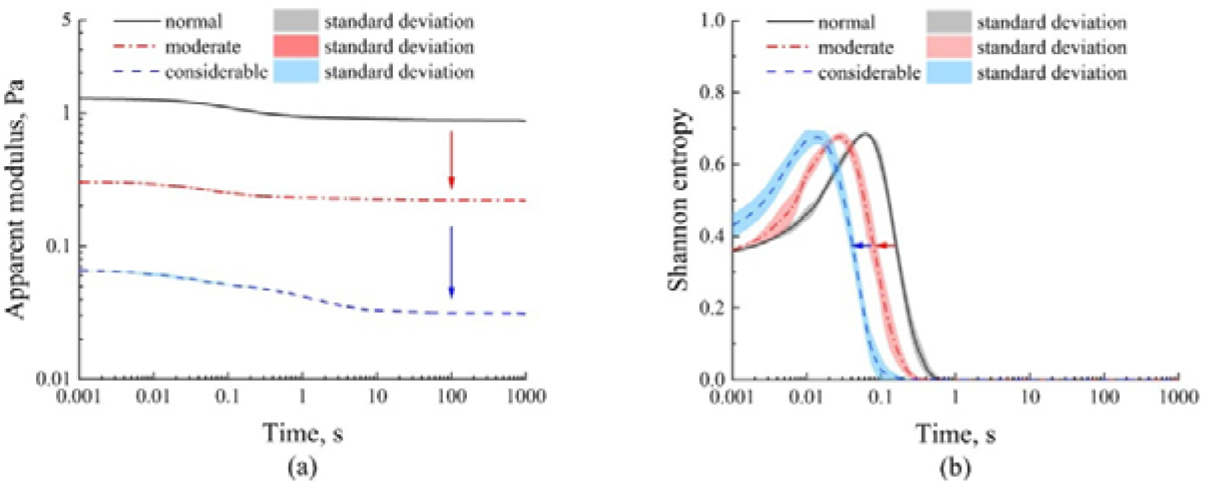
Response evolution of the cytoskeleton across different cancer progression stages, (a) apparent modulus, (b) Shannon entropy.

##### Evaluating drug intervention effects

To evaluate the efficacy of different drug treatments, we employed a single-parameter thought experiment. Starting from a considerable-softening state, we individually restored four key parameters—filament prestress, viscosity coefficient, filament density, and cross-linker dissociation rate—to a median level between the cancerous and normal cell states. The regulatory effects of these parameters on restoring the apparent modulus were analyzed, with the results presented in Fig. 6.

**Fig. 6.**
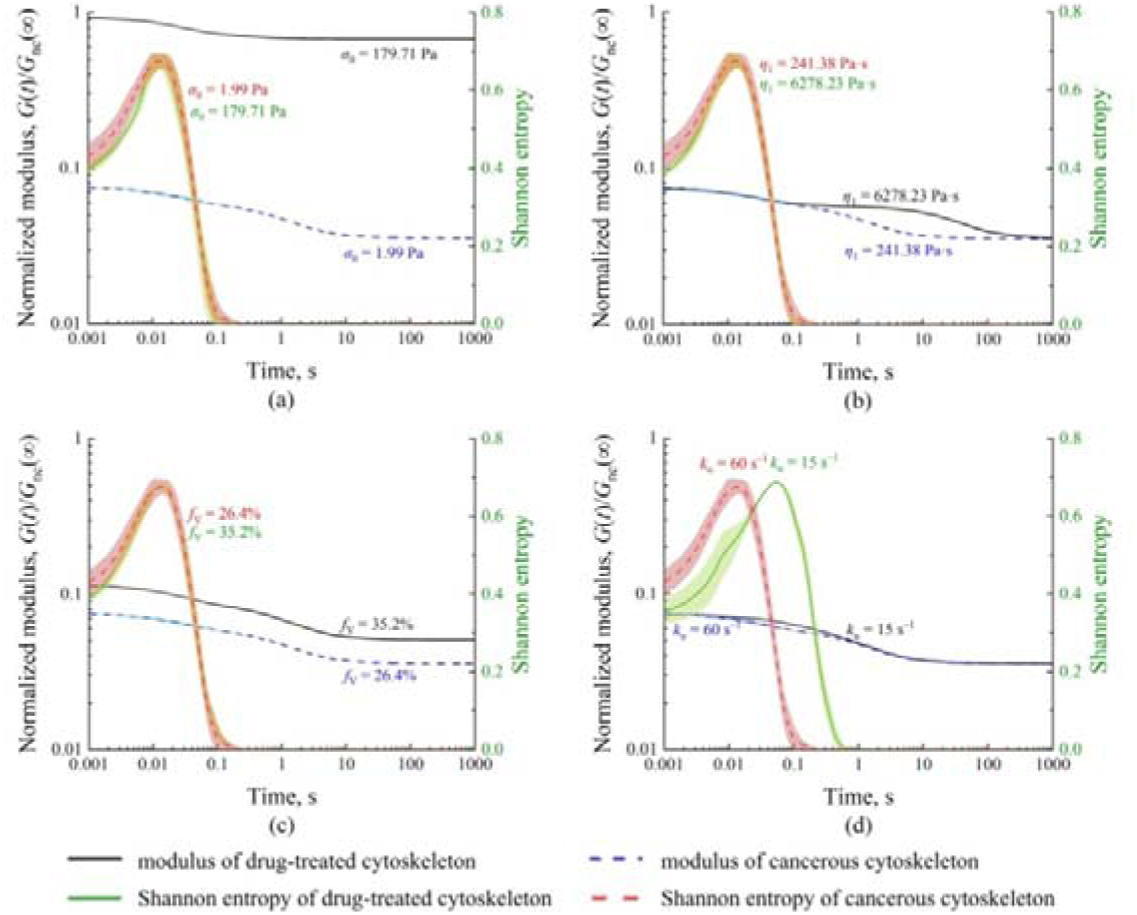
Effects of parameters related to cellular cancerous transform and drug treatment on the apparent modulus of cytoskeleton, (a) filament prestress, (b) filament viscosity parameter, (c) filament density, (d) cross-linker dissociation rate. (Blue dashed and black solid lines represent the modulus of cancerous and drug-treated cytoskeletons, respectively; red dashed and green solid lines represent the Shannon entropy of cancerous and drug-treated cytoskeletons, respectively).

## 3 Results

### 3.1 Cytoskeletal creep response

Fig. 3 shows the creep response of cytoskeletal networks with varying filament volume fractions under step shear stress. Within the short-term range of 0.001−0.7 s, the predicted response is close to the experimental measurements by Gardel et al. (2006) in both magnitude and power□law exponent. In the short- and intermediate-term ranges of 0.001–0.7 s and 0.7–11.57O s, the power□law exponent α gradually decreases with increasing filament volume fraction, and this trend is consistent with the FE simulations of Liang et al. (2023). Over longer times (*t* ≥ 11.57 s), the predicted response approaches a plateau stage, matching the long□term trend observed by Gardel et al. (2006).

### 3.2 Cytoskeletal time-dependent evolution

For a cytoskeletal network with a filament volume fraction of 35.2% under step shear stress, Fig. 4 shows its creep response evolution with four distinct temporal ranges, corresponding to the short-term (0.001–0.7 s, including first fast-varying and first transitional stages), intermediate-term (0.7–11.57 s), and long-term (*t* ≥ 11.57 s) ranges in Fig. 3.

In the first fast-varying stage (0.001 s ≤ *t* < 0.06 s), the bulk shear strain response (dark solid line in Fig. 4(a)) exhibits a positive power-law exponent, leading to fast response growth; the structural Shannon entropy (green solid line in Fig. 4(a)) rises, the contribution of bending filaments (red area in Fig. 4(b)) dominates its shear strain, and the count of cross-linkers shows a slow decline. In the first transitional stage (0.06 s ≤ *t* < 0.7 s), the response growth decelerates under a reduced power-law exponent; the Shannon entropy decreases, the contribution of filament deformation shifts toward a tension-dominated mode (purple area in Fig. 4(b)), and the count of cross-linkers undergoes a rapid decrease. In the intermediate-term range (0.7 s ≤ *t* < 11.57 s), the response growth becomes extremely slow; concurrently, the Shannon entropy remains near zero, the contribution of filament deformation becomes almost entirely tension-dominated, and the count of cross-linkers approaches near zero. Eventually, the response approaches a plateau in the long-term range (*t* ≥ 11.57 s).

### 3.3 Effects of cancerous transformation

Simulation results for the apparent shear modulus and Shannon entropy of the cytoskeleton at normal, moderate-softening, and considerable-softening cancerous cells are presented in Fig. 5. Compared to the normal cytoskeleton (black solid line), the moderate- and considerable-softening cytoskeletons (red dotted and blue dashed lines, respectively) not only soften significantly (Fig. 5(a)) but also complete network remodeling more rapidly (indicated by the leftward curve shift in Fig. 5(b)).

### 3.4 Effects of drug treatment on considerable-softening cancerous cytoskeleton

We systematically analyzed the impact of four key drug-treatment-related parameters —filament prestress, viscous coefficient, density, and cross-linker dissociation rate—on the normalized apparent modulus and structural Shannon entropy of considerable-softening cancerous cytoskeleton through a single-parameter thought experiment. The effects of individually tuning each parameter are illustrated in Fig. 6. Note that, *G*_nc_(∞) represents the steady-state apparent modulus (*t*→∞) of the normal cytoskeleton.

Comparing to the considerable-softening cancerous cytoskeleton, increasing filament prestress resulted in a consistently higher modulus throughout the simulated time (Fig. 6(a)); increasing the viscosity coefficient prolonged the time for the modulus to reach a plateau (Fig. 6(b)); increasing filament density provided a slight enhancement in the modulus (Fig. 6(c)); and reducing the dissociation rate resulted in a higher short-term modulus (*t* < 0.3 s) and delayed the peak in Shannon entropy from 0.016 s to 0.7 s (Fig. 6(d)).

## 4 Discussion

### 4.1 Model validation and capabilities

The multiscale cytoskeletal network model developed in this study demonstrates a strong capability to simulate the complex shear rheological behavior of cytoskeleton. As shown in Fig. 3, the model successfully reproduces the multiple power-law creep behavior observed in experimental shear rheology experiments (Gardel et al. 2006; Lieleg et al. 2010; Liang et al. 2023). Notably, it overcomes the temporal limitations associated with some traditional FE dynamical models (Saunders and Voth 2013), enabling the prediction of not only the short-and intermediate-term response of cytoskeleton but also its long-term, plateau-like creep response at extended timescales (>11.57 s) (Lieleg et al. 2010; Nam et al. 2016). Furthermore, the model demonstrates the capability to simulate cellular shear rheological behavior under pathological conditions, qualitatively replicating the experimental trend of "decreased modulus" and "advanced response" in cancerous cytoskeletons (Fig. 5, Fletcher and Mullins 2010; Hang et al. 2022; Xu et al. 2025).

It should be noted that the slight deviations between the predicted creep response and the experimental data of Gardel et al. (2006). (Fig. 3) may result from differences in material parameters. Our model parameters were taken from experiments on single stress fibers (Kumar et al. 2006) rather than from reconstituted FLNaOFOactin networks (Gardel et al. 2006). Additionally, the limited availability of specific data for node density and stiffness necessitated reliance on established cytoskeleton experimental or simulated values, which may collectively account for the observed discrepancies.

### 4.2 Mechanisms of power-law creep response

As shown in Fig. 4, the simulated cytoskeletal network exhibits a distinct four-stage creep response under step shear stress. These stages correlate with the evolving deformation state of the cytoskeletal filaments and the bio-chemo-mechanical properties of the cross-linkers, collectively driving continuous network remodeling. In the first fast-varying stage (0.001 s ≤ *t* < 0.06 s), cross-linked nodes impose strong rotational restraint on the filaments, promoting local filament bending deformation and increasing network Shannon entropy. This leads to a rapid rise in bulk strain with a positive power-law exponent. In the first transitional stage (0.06 s ≤ *t* < 0.7 s), the depolymerization of cross-linked nodes weakens their rotational constraint, shifting filament deformation toward a tension-dominated mode. Consequently, Shannon entropy decreases, and the bulk strain growth decelerates a reduced power-law exponent. In the intermediate-term range (0.7 s ≤ *t* < 11.57 s), local node connections stabilize, filament deformation becomes entirely tension-dominated, the Shannon entropy of the network stabilizes, and the bulk strain enters a slow creep process dominated by filament viscoelasticity; and eventually entering a plateau in the long-term range (*t* ≥ 11.57 s).

The present model captures the interactive coupling mechanism between node dynamics and filament deformation—a feature that is difficult to replicate using traditional affine network models or purely continuum approaches (Burla et al. 2019; Wang et al. 2021). By integrating the bio-chemo-mechanical properties of cross-linked and entangled nodes with filament viscoelasticity, our model reveals the microscopic origin for the multi-stage creep response, consistent with analytical evidence that characteristic relaxation/creep stages can be governed by intrinsic structural evolution factors coupled with deformation-state transitions (Cai et al. 2022). Moreover, the revealed network remodeling mechanism—governed by the coupling of node properties and filament deformation states—is not limited to cytoskeletal

networks. It may also be extended to other polymer network systems such as hydrogels and colloidal gels.

### 4.3 Regulation of drug treatment-related parameters

Our systematic analysis further elucidates the distinct mechanical roles of the four drug treatments-relate parameters in regulating the cancerous cytoskeletal behavior (Fig. 6). Both filament prestress and filament density can enhance the steady-state apparent shear modulus (Fig. 6(a) and 6(c)), with prestress being the key determinant. Similarly, both filament viscosity coefficient and cross-linker dissociation rate govern the processivity of apparent modulus evolution and network remodeling (Fig. 6(b) and 6(d)). Specifically, a reduced dissociation rate not only enhances short-term modulus but also significantly delays the peak time in structural Shannon entropy, indicating a prolonged remodeling process. These findings deepen the mechanistic understanding of cytoskeletal mechanical evolution during cancerization process and provide a theoretical basis for evaluating drug-treatment strategies. For example, while inhibiting cross-linker dissociation effectively delays network remodeling, synergistic interventions directly regulating filament prestress may still be required to restore the transient modulus effectively. This underscores the importance of combined therapeutic strategies in which the prognostic effects of modulating different parameters should be carefully evaluated.

We believe the extension of this model to polymer network materials will aid in simulating the power-law responses of the actin gel observed by Lieleg et al. (2010). It may also elucidate the mechanisms underlying network performance changes regulated by chemical cross-linkers, as discovered by Nam et al. (2016) in synthetic hydrogel. Thus, this model provides theoretical support for fields requiring co-regulation of nominal modulus and relaxation time, such as soft robotics and tissue engineering, and may facilitate the rational design of dynamic polymer networks.

### 4.4 Model limitations

It should be noted that despite its strong predictive capabilities, this model still has certain limitations. For instance, it does not account for the mobility or failure of physical entanglements between actin filaments (Liu et al. 2024), the active contraction units like myosin (Burla et al. 2019), the influence of heterogeneous structures like the cell nucleus (Oakes et al. 2017), or the three-dimensional network effects of the cytoskeleton (Wei et al. 2021). Furthermore, the model parameters rely heavily on limited experimental data (Kumar et al. 2006; Wei et al. 2021), and their generalization requires integration of more multi-scale material data; the current validation scope is also limited, necessitating tests across more cell types and pathological models (Hang et al. 2022). To address these limitations, future research should focus on introducing active mechanisms, expanding to three-dimensional models, and integrating machine learning for parameter identification. These approaches are essential to enhance the model’s predictive power and broader applicability.

## 5 Conclusion

This study establishes a multicomponent cytoskeletal network model that correlates, across multiscale, the bio-chemo-mechanical properties of cross-linkers, the viscoelasticity of actin filaments and their bending-to-tension transition, the dynamic remodeling of network structure, and the bulk shear rheological response. The model not only reveals the mechanism underlying the temporal evolution of cytoskeletal shear rheological properties, but also overcomes the temporal limitations of existing dynamic network models, enabling reasonable predictions of cytoskeletal shear rheological behavior over extended timescales. The key conclusions are as follows:

1. The synergistic and competitive interactions between the dissociation/association of cross-linkers and the transition of filament deformation states govern both the dynamic remodeling of cytoskeletal network and the multiple power-law characteristics of its bulk shear rheological response. Meanwhile, the inherent viscoelastic properties of the filaments determine the steady-state creep response of cytoskeletal network.
2. Filament prestress is a key parameter regulating the steady-state apparent modulus of cytoskeletal network. Effectively enhanced filament prestress serves as an effective strategy to reverse the abnormally decreased modulus caused by cancerous transforming. The process of cytoskeletal network remodeling is governed by the dissociation rate of cross-linked nodes, which together with the viscosity coefficient of filaments co-regulates the transient modulus.

The above insights provide a theoretical basis for understanding the mechanisms underlying the cancerous and drug-treated affections on cellular mechanical properties, and novel ideas for designing polymer networks that actively induce changes in cellular network structures. However, the predictive capability of the model remains constrained by the completeness of multiscale experimental data. Future work should integrate additional cytoskeletal components and active regulatory mechanisms.

## Author Contributions

Han-Lin Liu: Conceptualization (equal), Methodology (lead), Formal analysis (lead), Validation (lead); Visualization (lead), Writing - original draft (lead).

Neng-Hui Zhang: Conceptualization (lead), Writing - review & editing (equal), Supervision (lead), Funding acquisition (lead).

Jing-Jie You: Data curation (equal), Writing - review & editing (equal). Qi-Qi Li: Data curation (equal), Writing - review & editing (equal).

Cheng-Yin Zhang: Data curation (equal), Writing - review & editing (equal). All authors contributed to the manuscript and approved the submitted version.

## Statements and Declarations

The authors declare that they have no conflict of interest.

## Copyright

For Fig. 1(a), reprinted from Acta Biomaterialia, Vol 4(3), Jeffrey S. Palmer, Mary C. Boyce, Constitutive modeling of the stress-strain behavior of F-actin filament networks, Pages 597–612, Copyright (2008), with permission from Elsevier.

## Supplementary Information

The online version contains supplementary material available, providing finite element implementation details, component simplification, parameter determination, and advanced boundary treatments for the cytoskeletal network model.

## Data availability

All data that support the findings of this study are available on request from https://github.com/HanlinLiu-shu/Data-Cyto-Shearing.

## Supporting information

Supporting File: Solving Cytoskeletal Network Model Using Finite Element Method

## Acknowledgments

The authors thank Bo Gong for his useful suggestion on the refinement of the network model.

