## Supporting File: Solving Cytoskeletal Network Model Using Finite Element Method for "A multiscale cytoskeletal network model for shear rheological property and its evolutionary mechanism"

### Supporting File A: Solving Cytoskeletal Network Model Using Finite Element Method

This supporting file provides a comprehensive description of the finite element implementation for the cytoskeletal network model. The presentation follows the same structure as in the main text (Sections 2.2.1–2.2.3) and includes additional derivations and numerical procedures.

#### A1. Structural Discretization and Encoding

First, based on the non-uniform cytoskeletal network generated in Section 2.2.1, the network is divided into viscoelastic beam elements, and the nodes in the network are numbered sequentially from left to right and from bottom to top (e.g., the first row is numbered 1–*j*, the second row *j*+1–2*j*, and so on), resulting in a total number of nodes *n* = *i* × *j* (where *i* = 1, 2, 3, …, *N*_n_; *j* = 1, 2, 3, …, *N*_n_). Here *N*_n_ denotes the number of nodes per row. Each node is assigned three degrees of freedom (horizontal displacement *u^e^*, vertical displacement *v^e^*, and rotation *γ^e^*), leading to a total degree of freedom of 3*n*. Then, using node 1 as the origin, a global coordinate system is defined (horizontal as *X* and vertical as *Y*), with elements numbered first for vertical elements and then for vertical elements (for example, a horizontal element connects nodes (*m*, *m* + 1) and a vertical element connects nodes (*m*, *m* + *j*). A local coordinate system is established for each element (with the axial direction as *x^e^* and the transverse direction as *y^e^*, where *e* is the element number). Finally, considering the influence of time terms on node constraints, a post- processing method is employed. The global index for the free node *m* is set as (3*m* – 2, 3*m* – 1, 3*m*), and thus the position vector for element *e* connecting nodes *i* and *j* is given as

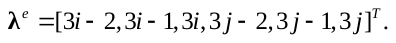
(A1)

#### A2. Element/Global Stiffness Matrix

Here, considering the constraining effect of node rotational springs, the stiffness of rotational springs is added to the corresponding diagonal elements of the local stiffness matrix. The specific steps are as follows: first, for a planar elastic beam, the element stiffness matrix in its local coordinate system is the summation of axial stiffness and bending stiffness, which takes the following form

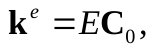
(A2)

where **C**_0_ is the parameter matrix obtained by extracting the material’s elastic modulus *E* from the traditional elastic element stiffness matrix **k**^e^ in the local coordinate system, and it satisfies

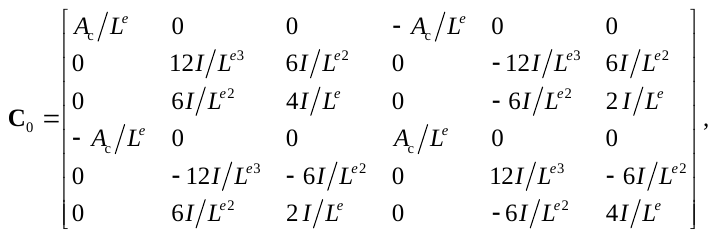
(A3)

where *A*_c_ is the cross-sectional area, *I* is the moment of inertia, and *L^e^* is the element length.

Considering the viscoelastic properties of cytoskeletal filaments, as described in section 2.1 (Eq. (2) in main text, and **Supporting File B1** for the filament viscoelastic parameters and prestress), it is necessary to modify the original stiffness matrix by replacing the modulus *E* with the time-dependent relaxation modulus *Y*(*t*) according to the principle of correspondence. This yields the corresponding time-dependent stiffness matrix, which is expressed as

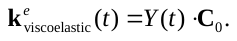
(A4)

Considering the distribution of physically entangled nodes and chemically cross-linked nodes described in Section 2.1, we took the spring stiffness *k*_disso_ (≈ 0) and *k*_cross_ (= *k*_0_) to generate the local spring stiffness matrix, respectively, namely

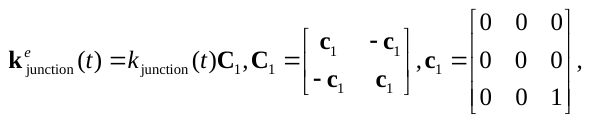
(A5)

where *k*_junction_(*t*) represents the rotational spring stiffness corresponding to the element node at time *t*, and it needs to be combined with the node state evolution equation (Eq. (3) in main text and the detailed description in **Supporting File B2**) to describe the dynamic changes of the nodes within the network.

Finally, considering the angle *θ^e^* between the cytoskeletal filaments in the global coordinate system, we performed a coordinate transformation on the local stiffness matrix to obtain the element stiffness matrix of cytoskeletal filaments in the global coordinate system, namely

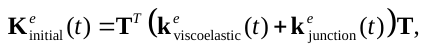
(A6)

where **T** is the coordinate transformation matrix, which satisfies

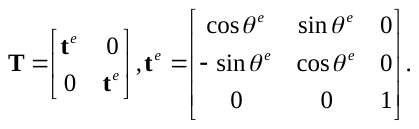
(A7)

Based on the numbering of nodal degrees of freedom at both ends of the element, the element **K***^e^*_initial_(*t*) is assembled into the corresponding positions of the global stiffness matrix. It is important to note that due to the introduction of time terms and the dynamic coupling of the nodes, the incremental method should be used. This involves discretizing time into time steps of Δ*t*, and updating the instantaneous elastic modulus *Y*(*t*), instantaneous nodal stiffness *k*_junction_ (*t*), and the total stiffness matrix of the network at each step.

#### A3. Generation of load arrays and handling of boundary conditions

Considering the spring constraints and the limiting effects of load-induced intracellular fluid on the cytoskeleton (approximated as a unit shearing stress *τH*(*t*), where *τ* = *τ*_0_/*W* (Iravani et al., 2020) on the outer network surface), an equivalent nodal load conversion is required. For uniformly distributed loads on the outer surface, they can be equivalently treated as a fixed-end force, i.e.

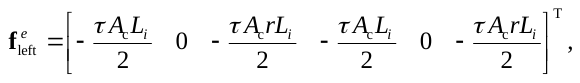
(A8)

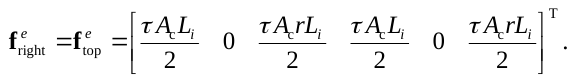
(A9)

Subsequently, the equivalent nodal forces from the local coordinate system are converted to the global coordinate system, and the load array is assembled based on the position vectors
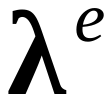
, and

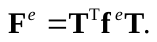
(A10)

Considering the coupling effect of external loads and the time factor on the performance of cytoskeletal filaments and the state of cross-linked nodes, we employed a time discretization method to divide the loading process into time steps of Δ*t*. This approach, together with the incremental iterative method, creates a nested solving structure for performing quasi-static analysis and displacement solving of cytoskeletal network.

Based on the equilibrium equations and boundary conditions, an iterative solution is performed. On one hand, a balance check is conducted to verify whether the nodal forces satisfy equilibrium at each time step. On the other hand, considering the restriction effect of intracellular fluid induced by cytoskeleton (Iravani et al., 2020), it is essential to verify that the governing equations (Eq. (5) in main text) are satisfied by the external boundary elements of cytoskeletal network at each time step. The detailed derivation of the governing equation and the advanced boundary treatments are provided in **Supporting File C**.

#### A4. Calculation, Verification, and Output

Considering the transition in local cytoskeletal filament deformation states (between bending- and tension-dominated modes) caused by filament viscoelasticity and node biochemical-mechanical characteristics, we employed a nested solution structure for quasi- static analysis and displacement determination of the cytoskeletal network. This is achieved through time discretization combined with an incremental iterative method.

In the time domain, to efficiently capture the responses from short-term (10^–2^ s) to long-term (10^–3^ s) scales, a segmented adaptive time-stepping strategy on a logarithmic scale is adopted. Specifically, the time axis is divided into segments bounded by points [10^–2^, 10^–1^, 0, 10^1^, 10^2^, 10^3^]. Within each segment, a fixed time step equal to 1/1000 of the segment’s end point is used. This strategy balances accuracy in the initial fast response with computational efficiency for long-term relaxation. With the current time being *t_n_*_+1_ = *t_n_* + Δ*t*. The modulus and external load at the current time step are updated according to Eqs. (2) and (A10), namely

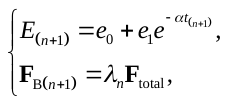
(A11)

where *λ_n_* is the load proportionality factor. Under the step loading conditions considered the step loading conditions considered here, *λ_n_* = *n*Δ*t*/*T* = 1. In the space domain, a nested iterative procedure is used. At each time step, the Newton-Raphson iterative method is employed until convergence is achieved. The specific collaborative solution steps are as follows.

**Step 1. Self-balance of initial state.** For *t* = 0^–^, the initialized node displacements are assigned based on experimental analysis, assuming the initial displacement conditions satisfy (Iravani et al., 2020)

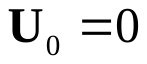
.	(A12)

Using the initial overall stiffness matrix (Eq. (4)) and the residual expression, we obtain

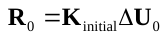
.	(A13)

The prestressed cytoskeleton exhibits self-balancing properties at *t* = 0⁻ (Kumar et al., 2006; Wei et al., 2020). To numerically determine this physical equilibrium state, we adopted the criterion of Iravani et al.(2020), deeming the network to be in initial equilibrium state when the residual norm for all nodes satisfies //**R**_0_^(^*^k^*^)^// ≤ 10^–6^.

**Step 2. Quasi-static equilibrium process.** For *t* ≥ 0^+^, the initial displacement condition is taken from the previous time step. The residual and tangent stiffness matrices are then expressed based on the reassembled overall stiffness matrix, namely

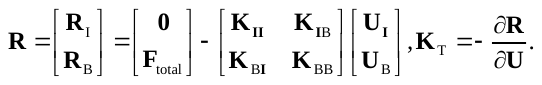
(A14)

Starting from iteration *k* = 0, we calculated the residuals and solved for the displacement increments, namely

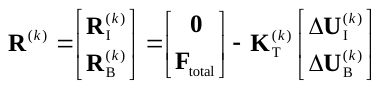
.	(A15)

Considering the viscoelastic deformation of filaments and node constraints, internal nodes are assumed to be in approximate equilibrium state at each time step. Thus, at current iteration *k*, both the residual norm conditions (//**R**^(^*^k^*^)^// ≤ 10^–6^) and the displacement convergence conditions (//**U**^(^*^k^*^)^// ≤ 10^–4^) should be satisfied (Iravani et al., 2020). Once the displacement correction **U**^(^*^i^*^+1)^ = **U**^(^*^i^*^)^ +Δ**U**^(^*^i^*^)^ meets these conditions, the converged displacement **U**_(_*_n_*_+1)_ is saved.

**Step 3. Judgment of cross-linked node states.** Based on Eq. (3), the node force, the dissociation and re-association rates (*k*_u_ and *k*_b_*N*_B_) for cross-linked nodes under the current deformation state are updated. Node state transitions are then evaluated by

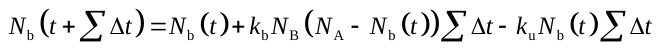
.	(A16)

Specifically, if |*N*_b_(*t* + ΣΔ*t*)–*N*_b_(*t*)| ≥1/*R*_cl_, where *R*_cl_ is the count of cross-linked nodes, an integer jump (i.e., cross-linker’s dissociation) is detected. It means that, one of the cross-linked nodes transitioned to the state of dissociated state. The rotational stiffness (Eq. (4)) of that node is changed from *k*_cross_ to *k*_disso_, and the element stiffness matrix is reset and then embedded into the overall stiffness matrix, incorporating the modified rotational cross-linked node changes. **Step 2** is repeated until the residual and displacement increment meet the tolerance requirements, i.e. |*k*_b_*N*_B_(*N*_A_–*N*_b_) –*k*_u_*N*_b_| <1%, after which the results at the current time step are output.

**Step 4. Post-processing and result output.** At each time step, a quasi-static solution for the cytoskeletal network is obtained via Newton-Raphson iteration, determining nodal displacement response. Statistical analysis of displacement patterns across all nodes characterizes the macroscopic deformation of cytoskeletal network at different time scales, enabling analysis of its shear rheological properties. Under a step shear stress *τ*_0_*H*(*t*), the shear strain creep response *γ*(*t*) and apparent shear modulus of the cytoskeletal network are given by (Efremov et al., 2020; Iravani et al., 2020)

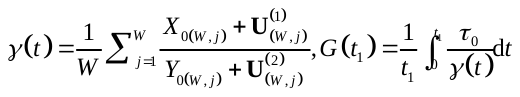
,	(A17)

where *j* is the column index of the cytoskeletal network, *X*_0(_*_W_*_，_*_j_*_)_ and *Y*_0(_*_W_*_，_*_j_*_)_ are the initial (*t_n_* = 0) global coordinates of the *j*-th boundary node; *τ*_0_ is the amplitude of unit shear stress, and *G* (*t*_1_) is the apparent shear modulus at time *t*_1_.

To further reveal the microstructural evolution, the deformation modes of all filaments at time *t*_1_— classified as bending- and tension-dominated modes, with proportions *p*_b_(*t*_1_) and *p*_s_(*t*_1_), respectively— are statistically analyzed. The Shannon entropy describing the bulk configuration change of the network is calculated as (Sajjadi et al., 2024)

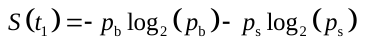
.	(A18)

For the calculation of network deformation, the buckling state of compressed cytoskeletal filaments should also be assessed (Yu et al., 2025). As shown in Fig. A1(a), after conducting a mesh independence verification, we selected a cytoskeletal network composed of 11×11 units for simulation. During the global calculation process, we observed the changes in results by reducing Δ*t* to verify the stability of the computational results and to determine whether to output them, ensuring the independence of the time step. As shown in Fig. A1(b), the computational results in this study have met the stability conditions.

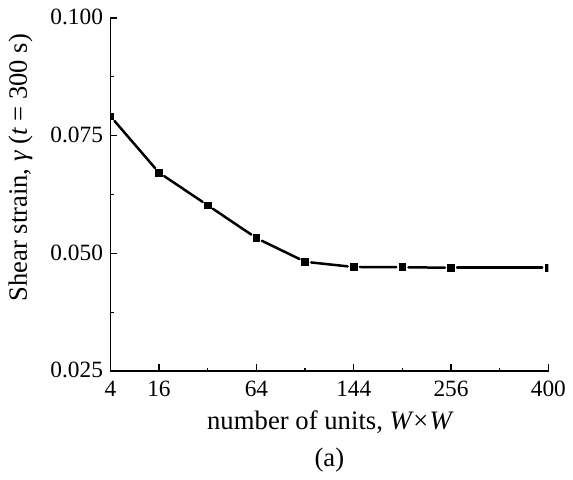

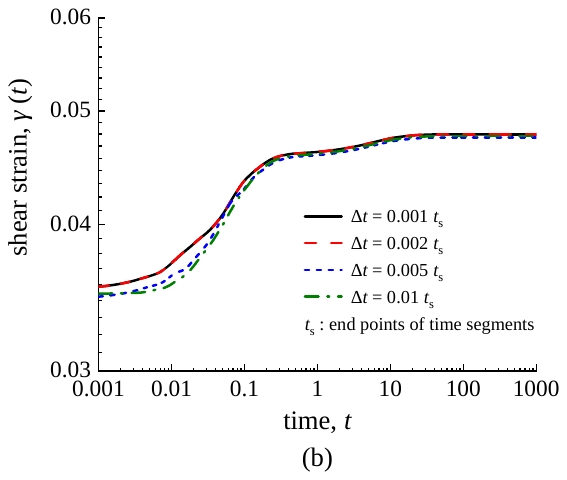

Fig. A1. Convergence verification of FE simulation: (a) mesh independence verification, (b) time stability verification.

### Supporting File B: Component Simplification and Parameter Determination

#### B1. Node-Scale Simplification and Bio-chemo-mechanical Modeling

As illustrated in Fig. 1(c) (main text), the linked nodes exhibit diverse physical or chemical bonding mechanisms (Kurniawan et al.,2012; Broedersz et al., 2014; Liu et al., 2024). Therefore, at the node scale, based on experimentally derived biochemical-mechanical properties, we classified them into physically entangled nodes connected by weak interactions and chemically cross-linked nodes connected by specific cross-linkers (Kurniawan et al., 2012; Broedersz et al., 2014). Due to their weak interactions, the physically entangled nodes are simplified as hinge nodes that transmit force but not bending moment (Kurniawan et al., 2012), which are assumed not to fail. Chemically cross-linked nodes not only constrain the rotation of connected filaments (Zhang et al., 2021) but also exhibit force-regulated dissociation/association rates, following the Bell model (Bell, 1978). These cross-linked nodes are characterized using a biochemical-mechanical coupling approach that simultaneously describes their dissociation/association behavior and constraint effects on filaments.

On the one hand, the rate equation derived from the Bell model (Bell, 1978; Nam et al., 2016) describes the mechanically regulated dissociation/association of cross-linkers (i.e., changes in binding density), expressed as (Nam et al., 2016)

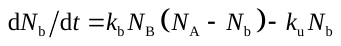
,	(B1)

where *N*_b_, *N*_A_ and *N*_B_ are the binding density, ligand density and receptor density of cross-linkers, respectively; *k*_u_ and *k*_b_*N*_B_ are the dissociation and re-association rates of cross-linkers, respectively, which follow the loading-regulated functions (Nam et al., 2016; Wei et al., 2021), i.e.

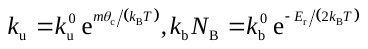
,	(B2)

where *k*_B_ and *T* are the Boltzmann constant and absolute temperature, respectively; *k*_u_^0^ and *k*_b_^0^ are the intrinsic dissociation and re-association rates under zero load, determined experimentally (Nam et al., 2016; Wei et al., 2021); *m* is the torque applied on a single cross-linked node, *θ*_c_ is the equivalent rotation angle for the node’s characteristic transition, and *E*_r_ represents the elastic energy stored in the re-formed cross-linker (Wei et al., 2021). And the Bell model parameters used in this study are summarized in Table 3 (main text).

On the other hand, the chemical cross-linked nodes are simplified as rotational spring nodes with stiffness *k*_j_, accounting for their transition from cross-linked to pre-cross-linked states during cross-linker dissociation and the concomitant weakening of their rotational constraint on filaments (Zhang et al., 2021). We utilize the rotational spring stiffness *k*_0_, which is related to the filaments volume fraction and the dynamic force properties of cross-linkers under zero stiffness conditions, as obtained by Zhang et al. (2021) through experiments and FE simulations. Owing to the lack of direct experimental measurements for the torsional stiffness of cross-linked nodes, we assumed that the relationship between the rotational spring stiffness *k*_0_ and the filament volume fraction *f*_V_ follows the model of Zhang et al. (2021) obtained through a combination of experiments and FE simulations, namely

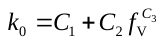
.	(B3)

where *C*_1_ = 1.07×10^–5^ N·μm/rad, *C*_2_ = 7.13×10^–2^ N·μm/rad, and *C*_3_ = 3.51.

In addition, FM experiments of the cytoskeleton have revealed that the ratio of physically entangled nodes to chemically cross-linked nodes exhibits an approximately linear trend across cells at different growth stages (Kurniawan et al., 2012). Therefore, we hypothesize that the count of entangled nodes *R*_e_ and cross-linked nodes *R*_cl_ follow the statistical relationship *R*_e_ = *s R*_cl_, where *s* is the proportionality constant.

#### B2. Filament-Scale Viscoelastic Parameter Determination from Laser Nanoscissor Experiments

The data on viscoelastic properties of cytoskeletal filament is scarce. Therefore, we used laser nano-scissor experimental data on stress fibers from Kumar et al. (2006) (Fig. B1) to determine the prestress and viscoelastic parameters for our model of cytoskeletal filaments (Wei et al., 2021; Liang et al., 2023). Our analysis of this experimental data (Kumar et al., 2006) revealed that the instantaneous retraction length of the fibers significantly exceeded the diameter of the ablated region and was influenced by different drug treatments. Furthermore, the instantaneous retraction length, regulated by the prestressed state, also exhibits various changes (Gardel et al., 2006; Wei et al., 2021). We therefore hypothesize that the instantaneous retraction results from both laser ablation and the transient elastic recovery of the fibers, as shown in Fig. B1(a). Consequently, a three-parameter solid model (Lyu et al., 2020) was selected to describe the viscoelasticity of stress fibers, and the experimental results were processed accordingly.

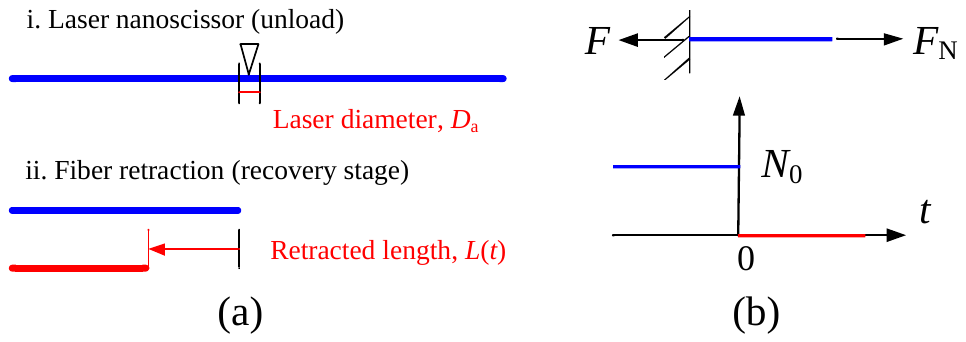

Fig. B1. Schematic of laser nano-scissor experiment on a single intracellular stress fiber, (a) simplified representation of stress fiber retraction after laser nano-scissoring, (b) force applied to stress fiber.

First, the in vivo laser nano-scissor experiment was simplified. As shown in Fig. B1(b), prior to laser nano-scissoring, the end of stress fiber experiences an axial pretension force *N*_0_ related to cell tension, environmental load, and fiber-fiber interactions (Onck et al., 2005; Gardel et al., 2006; Wei et al., 2021). Fiber breakage signifies the unloading of this prestress. Therefore, the loading protocol in the laser nano-scissor experiment is approximated as

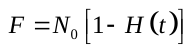
,	(B4)

where *N*_0_ = *σ*_0_ *A*, in which *σ*_0_, *A* and *F* are the prestress, cross-sectional area and the change in tensile force of stress fiber, respectively; *H*(*t*) is the unit step function. Substituting Eqs. (2) and (B4) into the force balance equation (*F* = *F*_N_) yields

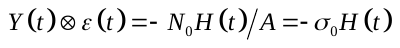
.	(B5)

Expanding Eq. (B5) in the form of Eq. (2) (Lyu et al., 2020), the fiber deformation accounting for the history effect is given as

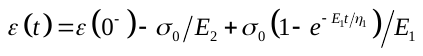
.	(B6)

Subsequently, the original and maximum lengths of stress fiber (*L*_F_ and *L*_max_) before laser nano-scissoring following drug treatment were obtained from experimental video (Table B1).

Table B1. Lengths of stress fibers under different drug treatments (Kumar et al., 2006)

| Drug treatment | *L*_s_ (μm) | *L*_max_ (μm) |
| --- | --- | --- |
| Drug-free | 6.64 | 10.00 |
| ROCK | 6.64 | 8.54 |
| MLCK | 6.64 | 6.81 |

Referring to Table B1, the time-history curve of fiber retraction length was converted to that of fiber deformation. The prestrain *ε*(0^–^) before laser nano-scissoring (*t* = 0^–^) and the instantaneous strain *ε*(0^+^) after scissoring (*t* = 0^+^) are given as

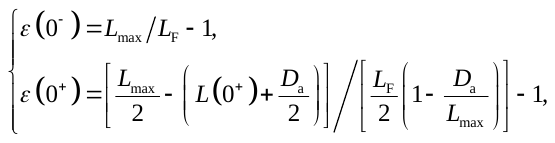
(B7)

where *L*(0^+^) is the instantaneous retraction length after laser nano-scissoring (Kumar et al., 2006). Substituting the instantaneous strains before and after fiber breakage into Eq. (B6), and combining with *ε*(0^+^) = *ε*(0^–^)–*σ*_0_/*E*_2_, we obtained the relationship between the transient modulus *E*_2_ and the prestress *σ*_0_, reducing the number of fitting parameters.

Finally, the parameter fitting space was defined based on the established ranges from literatures. Specifically, the elastic modulus (*E*_2_ = 100~12000 Pa, Gardel et al., 2010; Liang et al., 2023), prestress of single actin filament (*σ*_0_ = 0~180 Pa, Kumar et al., 2006; Wei et al., 2021), and the viscosity parameter from filament-intracellular fluid interactions (*η*_1_ = 3.0~10000 Pa·s, Nam et al., 2016; Chang et al., 2024). Fitting the time-dependent deformation curves (Kumar et al., 2006) using Eq. (B6) yielded the filament viscoelastic parameters and prestress listed in Table 2 (main text).

### Supporting File C: Advanced Boundary Treatments

#### C1. Determination of quasi-static governing equation for external boundary cytoskeletal filaments using the virtual work principle

Based on the corresponding models selected in Section 2.1 to describe different scale components of cytoskeleton, we used the microelement method to determine the quasi-static governing equation for the external boundary cytoskeletal filaments.

Under the action of shear stress, the cytoskeletal network undergoes shear deformation. Considering the restriction effect of intracellular fluid induced by cytoskeleton, it can be assumed that the lower boundary of cytoskeletal network is fixed, while the remaining boundaries are subjected to an eccentric tangential uniform force *τH*(*t*)*A*_c_ (Iravani et al., 2020). Given that the boundary nodes are modeled as rotational spring hinge nodes holding a stiffness of *k*_j_ and geometric continuity conditions at the nodes, we extracted each cytoskeletal filament from the boundary of network individually. We established a local coordinate system as shown in Fig. C1(a), where the *x*-axis lies along the mid-surface of skeletal filament, the *y*-axis points outward from the cytoskeletal filament surface, and the lengths and radii of the cytoskeletal filaments are denoted as *L_i_* and *r*_F_, respectively. The outer surface of the cytoskeletal filament is assumed to be acted upon by the eccentric surface tangential uniform load *τA*_c_.

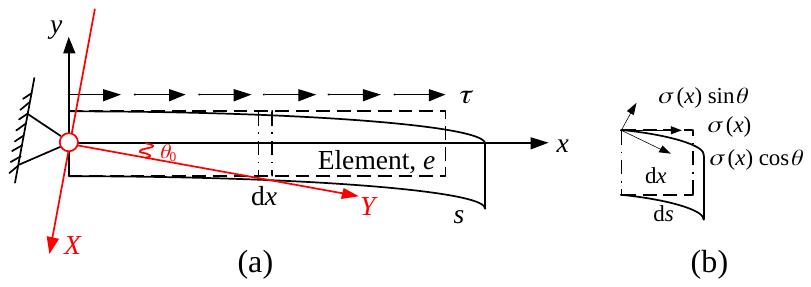

Fig. C1. Cytoskeletal filaments on the outer boundary of cytoskeletal network: (a) schematic diagram of filament deformation under force, (b) schematic diagram of deformation of a micro-element segment of the filament.

First, consider the work done by the surface forces. Under the eccentric tangential uniform surface stress shown in Fig. C1(a), using the method of sections, each cross-section of cytoskeletal fiber is subjected to an axial force *σ*(*x*) (= *τA_c_* (*L_i_* –*x*)). The work done by the surface stress can be divided into two parts: the tensile effect on the micro-beam caused by the component *σ*(*x*)cos*θ* in the tangent direction of deflection curve, and the work done by the rotational effect caused by the component *σ*(*x*)sin*θ* in the normal direction. Let *u_i_* be the displacement of a point along the *x* (or *s*) coordinate line in the *x* (or *s*) direction, and *w_i_* be the displacement of a point along the *y* (or perpendicular to *s*) coordinate line in the *y* (or perpendicular to *s*) direction. Let *κ* be the curvature after the cytoskeletal filament undergoes bending deformation. Under the small deformation assumption (i.e., the angle of rotation *θ* is small), based on the deformation of an arbitrary micro-element segment of cytoskeletal filament shown in Fig. B1(b), the deformations can be approximated as follows (Holzapfel and Ogden, 2011): *u*'(*s*) = d*u*(*s*)/d*s* = cos*θ*, *w*'(*s*) = d*w*(*s*)/d*s* = sin*θ* ≈ *θ*, d*w*/d*x* = *w*'(*s*)/*x*'(*s*) ≈ *w*'(*s*), *κ* ≈ *w*''(*s*) ≈ *w*''(*x*), (1+*u*'(*x*)) d*x* = cos*θ* d*s* ≈ d*s*. Therefore, the work done by the eccentric uniform tangential surface stress can be approximately expressed as

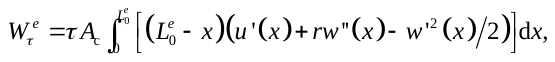
(C1)

where
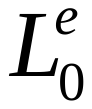
 is the length of cytoskeletal filament in the quadrilateral cytoskeletal unit, and *e* represents the filament index.

Considering the restriction of cross-linked nodes on filament rotation, the potential energy
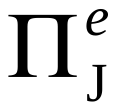
 of single rotational spring can be expressed as

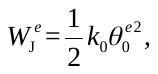
(C2)

where
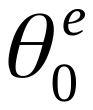
 is the initial shear angle of the filament after pre-stretching (Wei et al., 2021) . It is important to note that in this study, the physically entangled nodes are simplified to hinge nodes (Kurniawan et al., 2012), with a node stiffness of zero (*k*_0_ = 0). And the entanglements can also be described using Eq. (A5).

When the cytoskeleton undergoes local deformation, the energy of each individual cytoskeletal filament is composed of the stretching/compression strain energy
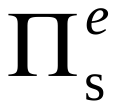
 and the bending strain energy
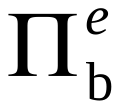
 (Holzapfel and Ogden, 2011). Therefore, the total energy
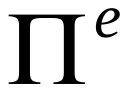
 of a single cytoskeletal filament can be expressed as

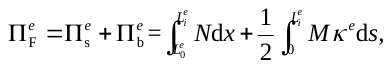
(C3)

where the axial force and bending moment can be written as
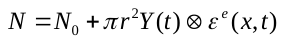
and
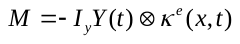
, respectively. And *N*_0_ is the pre-tension force of filament, *I_y_* is the moment of inertia of filament about the *y*-axis; 
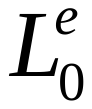
 is the length of filament with zero tensile/compressive deformation, and
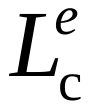
 is the contour length of filament (Holzapfel and Ogden, 2011).

By substituting the deformation approximation based on the small deformation assumption into Eq. (C3), the total energy of a single filament connected by cross-linked points can be expressed as

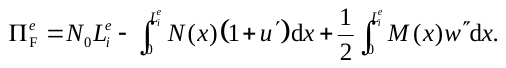
(C4)

By combining Eqs. (C1) and (C2), the virtual work of external forces and the virtual work of deformation can be described using the virtual displacement, leading to

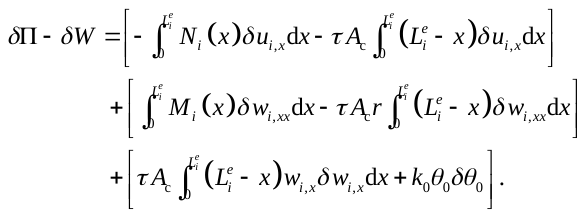
(C5)

By applying integration by parts to Eq. (C5), we have

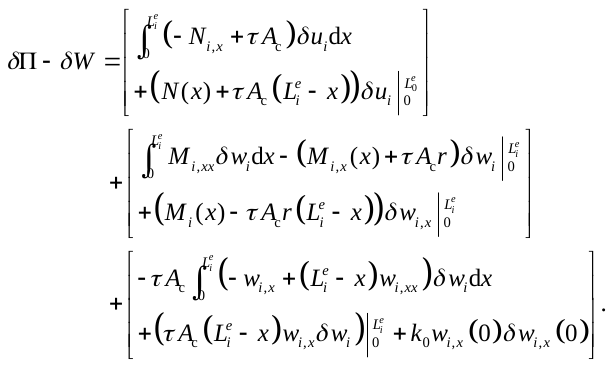
(C6)

Using the principle of virtual work
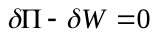
 and the arbitrariness of each virtual displacement, the quasi-static governing equation for cytoskeletal filament under the action of eccentric uniformly distributed tangential surface stresses can be expressed as

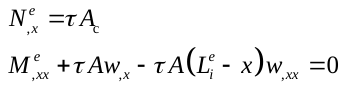
(C7)

where *N^e^* and *M^e^* are the axial force and bending moment of filament element *e* at the current time step *t_n_* = *n* + 1, respectively. Their discrete forms satisfy

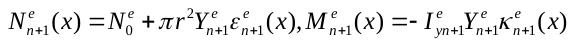
.	(C8)

#### C2. Stiffness Matrix Correction for Boundary Elements

At the current time step, the internal nodes (subscript I) and boundary nodes (subscript B), together with the overall stiffness matrix, displacement vector, and load vector, are represented in block form as

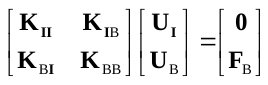
,	(C9)

where **K**_II_ is the sub-stiffness matrix for elements with both ends as internal nodes; **K**_IB_ and **K**_BI_ are the coupling sub-stiffness matrices between internal and boundary node degrees of freedom; and **K**_BB_ is the sub-stiffness matrix for elements with both ends as boundary nodes, which requires geometric correction. **U**_I_ and **U**_B_ are the displacements of internal and boundary nodes, respectively, and **F**_B_ is the force applied to the boundary nodes.

For the internal stiffness matrix **K**_II_, only the direct coupling of stiffness between the degrees of freedom for internal nodes needs to be considered, we have

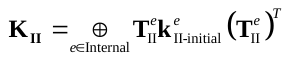
,	(C10)

where
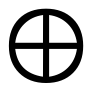
 denotes the assembly of element local stiffness matrices into the overall stiffness matrix in the global coordinate system; **T**_II_ is the coordinate transformation matrix for internal nodes; **k***^e^*_II-initial_ is the element stiffness matrix of internal filaments in the local coordinate system at the current time step, obtained from Eq. (4).

The boundary stiffness matrix **K**_BB_ comprises both the original local stiffness matrix **k*^e^*_BB_** of cytoskeletal filaments at the network and a correction term based on the governing equation. For axial boundaries, the tensile stiffness matrix **k*^e^*_BB-initial-str_** combined with the load array satisfies the discrete form of Eq. (C7). For lateral boundaries, however, the bending stiffness matrix **k*^e^*_BB-initial-bend_** combined with the load array satisfies only the linear part of Eq. (C7). To account for the influence of eccentric uniformly distributed loading on lateral bending, a geometric stiffness matrix is introduced as

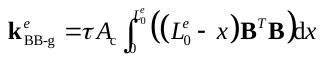
,	(C11)

where **B** is the curvature-displacement matrix, and **B***^e^* = ▽^2^**N***^e^*; **N** is the shape function. Combining the axial, bending, and geometric stiffness components yields the boundary- corrected element stiffness matrix in the local coordinate system and the boundary stiffness matrix in the global coordinate system, respectively, as

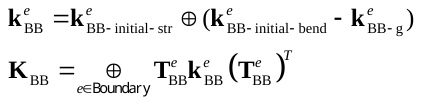
(C12)

where **T***^e^*_BB_ is the coordinate transformation matrix for boundary nodes; **k***^e^*_BB-initial-str_ and **k***^e^*_BB-initial-bend_ are the tensile and bending components, respectively, of the element stiffness matrix for boundary filaments in the local coordinate system at the current time step, obtained from Eq. (4). For cytoskeletal filaments (coupled elements) connecting internal and boundary nodes, their local coupling stiffness matrices **k**_IB_ and **k**_BI_ are expressed as

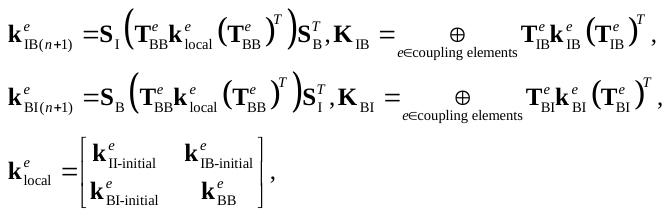
(C13)

where **k***^e^*_IB-initial_ and **k***^e^*_BI-initial_ are the local stiffness matrices of the coupled elements at the current time step, obtained from Eq. (4); **S**_I_ and **S**_B_ are the degree-of-freedom extraction matrices. Finally, embedding the modified element stiffness matrices into the overall stiffness matrix yields the boundary-modified overall stiffness matrix.

Furthermore, as described in **Supporting File A4**, a post-processing method is employed to handle boundary conditions, requiring adjustments to the constrained degrees of freedom. Considering the constraints observed on the upper and lower boundaries of the cell/cytoskeleton in shear rheological experiments and numerical simulations (Gardel et al., 2006), for the case of the fixed lower boundary, we apply a “1” method; for the vertical constraints on the upper boundary, we only constrain its vertical translational degrees of freedom; and for internal physical entangled (hinged) nodes, we set the corresponding rotational degrees of freedom and the respective rows/columns to zero.
